# The influence of familiarity on the neural coding of face sex

**DOI:** 10.1101/2022.10.27.514076

**Authors:** Celia Foster, Johannes Schultz, Melissa Munzing, Isabelle Bülthoff, Regine Armann

**Affiliations:** Max Planck Institute for Biological Cybernetics, Tübingen, Germany; Biopsychology and Cognitive Neuroscience, Faculty of Psychology and Sports Science, Bielefeld University, Germany; Center for Economics and Neurosciences, University of Bonn, Germany; Institute for Experimental Epileptology and Cognition Research, Medical Faculty, University of Bonn, Germany

**Keywords:** sex, gender, categorization, fMRI, OFA, FFA

## Abstract

In behaviour, humans have been shown to represent the sex of faces categorically when the faces are familiar to them. This leads to them judging faces crossing the category boundary (i.e. from male to female) as more different than faces that are within the same category. In this study, we investigated how faces of different sexes are encoded in the brain, and how familiarity changes the neural coding of sex. We recorded participants’ brain activity using fMRI while they viewed both familiar and unfamiliar faces that were morphed in their sex characteristics (i.e. between male and female). Participants viewed pairs of faces that were either identical, or differed in their sex morph level, with or without a categorical change in perceived sex (i.e. crossing the perceived male/female category boundary). This allowed us to disentangle physical and categorical neural coding of face sex, and to investigate if neural coding of face categories was enhanced by face familiarity. Our results show that the sex of familiar, but not unfamiliar, faces was encoded categorically in the medial prefrontal and orbitofrontal cortex as well as in the right intraparietal sulcus. In contrast, the fusiform face area showed a sensitivity to the physical changes in the sex of faces that was unaffected by face familiarity. The occipital face area showed its highest responses to faces towards the ends of the sex morph continuum (i.e. the most male or most female faces), and these responses were also unaffected by face familiarity. These results suggest that there is a dissociation between the brain regions encoding physical and categorical representations of face sex, with occipital and fusiform face regions encoding physical face sex properties and frontal and parietal regions encoding high-level categorical face sex representations that are linked to face identity.

## 1. Introduction

We can automatically determine the sex or gender of a face, as well as many other characteristics, even when we only view the face briefly. This ability is one example of the astounding capabilities of our visual face perception system, which can detect many characteristics (e.g. sex, identity, expression, race, age) despite a great variance in low-level visual properties of the faces we see (e.g. due to differences in lighting or viewpoint). Neuroimaging studies have identified several distinct brain regions that show strong responses when participants view images of faces (for reviews, see Haxby et al., 2000; Tsao & Livingstone, 2008). It is thought that this face network may be specialized to detect and process specific face characteristics, allowing us to robustly detect these face characteristics from varied low-level visual input.

Neuroimaging studies have investigated how face-responsive brain regions encode information about the sex of faces. Natural images of male and female faces were found to evoke different patterns of activity in several regions in the face-responsive network including the fusiform face area (FFA), occipital face area (OFA), superior temporal sulcus, inferior frontal gyrus, insula cortex and orbitofrontal cortex (OFC) (Contreras et al., 2013; Kaul et al., 2011—but see Foster et al., 2019, who could not decode the sex of faces in these regions using face stimuli controlled for low-level visual properties). A study using faces morphed in sex found that the FFA responded to these morphs linearly, in alignment with the changes induced by the morphing procedure, whereas the OFC responded non-linearly, in alignment with participants’ behavioural judgments of the faces’ sex (Freeman et al., 2010). These studies show that many regions in the face network encode information about face sex, and that different regions may encode different face sex properties.

Behavioural studies have also helped reveal how face characteristics are encoded in the brain. It has been found that stimuli crossing a boundary between two categories can be perceived as more different than stimuli that do not, a phenomenon known as categorical perception (Harnad, 1987; Rosch et al., 1976). For faces, categorical perception has been found for several face traits including face identity (Beale & Keil, 1995), expression (Calder et al., 1996) and race (Levin & Angelone, 2002). For sex, studies using face stimuli that vary linearly in sex information have demonstrated categorical perception of sex for familiar faces, but not for unfamiliar faces (Armann & Bülthoff, 2012; Bülthoff & Newell, 2004). One earlier study that had reported categorical perception of sex for unfamiliar faces used stimuli obtained by morphing between male and female faces of different identities, opening the possibility that the effect was driven by categorical perception of face identity (Campanella et al., 2001). Overall, these studies suggest that familiarity induces categorical perception of face sex.

As previous neuroimaging studies have only investigated the neural coding of the sex of unfamiliar faces (Contreras et al., 2013; Foster et al., 2019; Freeman et al., 2010; Kaul et al., 2011), it is not yet known how the brain encodes the sex of familiar faces. In particular, it is unknown whether the neural coding of face sex becomes categorical with face familiarity, in line with behavioural evidence (Armann & Bülthoff, 2012; Bülthoff & Newell, 2004). A number of previous studies have demonstrated that neural responses in the FFA and anterior temporal face area (ATFA) are enhanced when participants recognise familiar faces (Grill-Spector et al., 2004; Hoffman & Haxby, 2000; Nasr & Tootell, 2012). Furthermore, several distributed regions in the face-responsive network have also been shown to encode face identity information (Andrews & Ewbank, 2004; Anzellotti et al., 2014; Anzellotti & Caramazza, 2016; Foster et al., 2021; Gauthier et al., 2000; Guntupalli et al., 2017; Jeong & Xu, 2016; Kriegeskorte et al., 2007; Loffler et al., 2005; Nestor et al., 2011; Winston et al., 2004). A study that specifically investigated which regions encode categorical and physical identity information for familiar faces found that the FFA encoded face identity in a categorical manner, whereas the OFA encoded physical aspects of face identity (Rotshtein et al., 2005). This finding suggests that physical and categorical aspects of face characteristics are encoded by different brain regions.

In the present study, we investigated which brain regions encode physical and categorical aspects of face sex for both familiar and unfamiliar faces. We created a stimulus set of 20 face identities that were morphed along the sex continuum but were unchanged in other non-sex related characteristics. We familiarised participants with half of the 20 face identities in the dataset. These participants then took part in an fMRI experiment, where we recorded their brain activity as they viewed both the unfamiliar and familiar faces. We used a repetition suppression design where participants viewed pairs of faces from three different conditions; (1) identical faces, (2) faces that varied in their sex morph level but did not cross the male/female category boundary, and (3) faces that varied to the same degree in their sex morph level and also crossed the male/female category boundary. These three conditions allowed us to disentangle neural coding of physical changes in face sex (i.e. higher responses to pairs of faces that varied in their morph level compared to identical faces) and to categorical changes in face sex (i.e. higher responses to pairs of faces that crossed the sex category boundary compared to faces that did not). To address our research question, participants viewed both unfamiliar and familiar faces, which allowed us to investigate whether the neural coding of face sex is altered by face familiarity.

## 2. Materials and methods

### 2.1. Participants

Twelve female participants (18–37 years old, mean = 24.5) completed the fMRI experiment. A previous behavioural study identified an effect size of Cohen’s *d* = 1.85 for higher maximum performance of face sex discrimination for familiar as compared to unfamiliar faces (Armann & Bülthoff, 2012). A power sensitivity analysis performed using the G∗Power3 software (Faul et al., 2007) indicated that a sample size of 10 would be required to detect this effect size at the 0.05 alpha level with 80% power. We chose to include only participants of one sex, as differences have been found between male and female participants’ gaze behaviour when they judge the sex of faces (Armann & Bülthoff, 2009). We thus avoided biases that could be caused by such variations in gaze behaviour by only including female participants. All participants provided written informed consent prior to the experiment, and the experiment was approved by the local ethics committee of the University Hospital Tübingen.

### 2.2. Stimuli

#### 2.2.1. Main experiment stimuli

Our experimental stimuli were taken from a dataset that was created and perceptually validated in a previous study (Armann & Bülthoff, 2012). This dataset consisted of 3D face scans of 20 individuals (10 male, 10 female) from the face database of the Max Planck Institute for Biological Cybernetics (https://faces.kyb.tuebingen.mpg.de/), that were aligned to a 3D morphable model (Blanz & Vetter, 1999) and morphed to create a same-sex to opposite-sex continuum for each identity by applying a sex vector consisting of the difference between an average male and an average female face. For the present study, we selected four morph levels from each identity’s sex continuum: 0% female (i.e. 100% male), 40% female, 80% female and 120% female (see **Fig. 1A**). We extended the female endpoint of the sex continuum (i.e. 120% female rather than 100% female) as it has been previously shown that observers have a bias towards perceiving faces as male, at least when tested with stimuli such as ours (Armann & Bülthoff, 2012; Luther et al., 2021). Thus, our four selected morph levels allowed for the point of subjective equality (change in perception of face sex) to lie between the 40% female and 80% female morph levels (Armann & Bülthoff, 2012). A further sex continuum was created for one additional identity, which was used as a target identity for the behavioural task performed during the fMRI experiment (see Section 2.3.2.). Faces were presented in grayscale and were oriented 20° to the right to aid face shape visibility.

**Figure 1.**
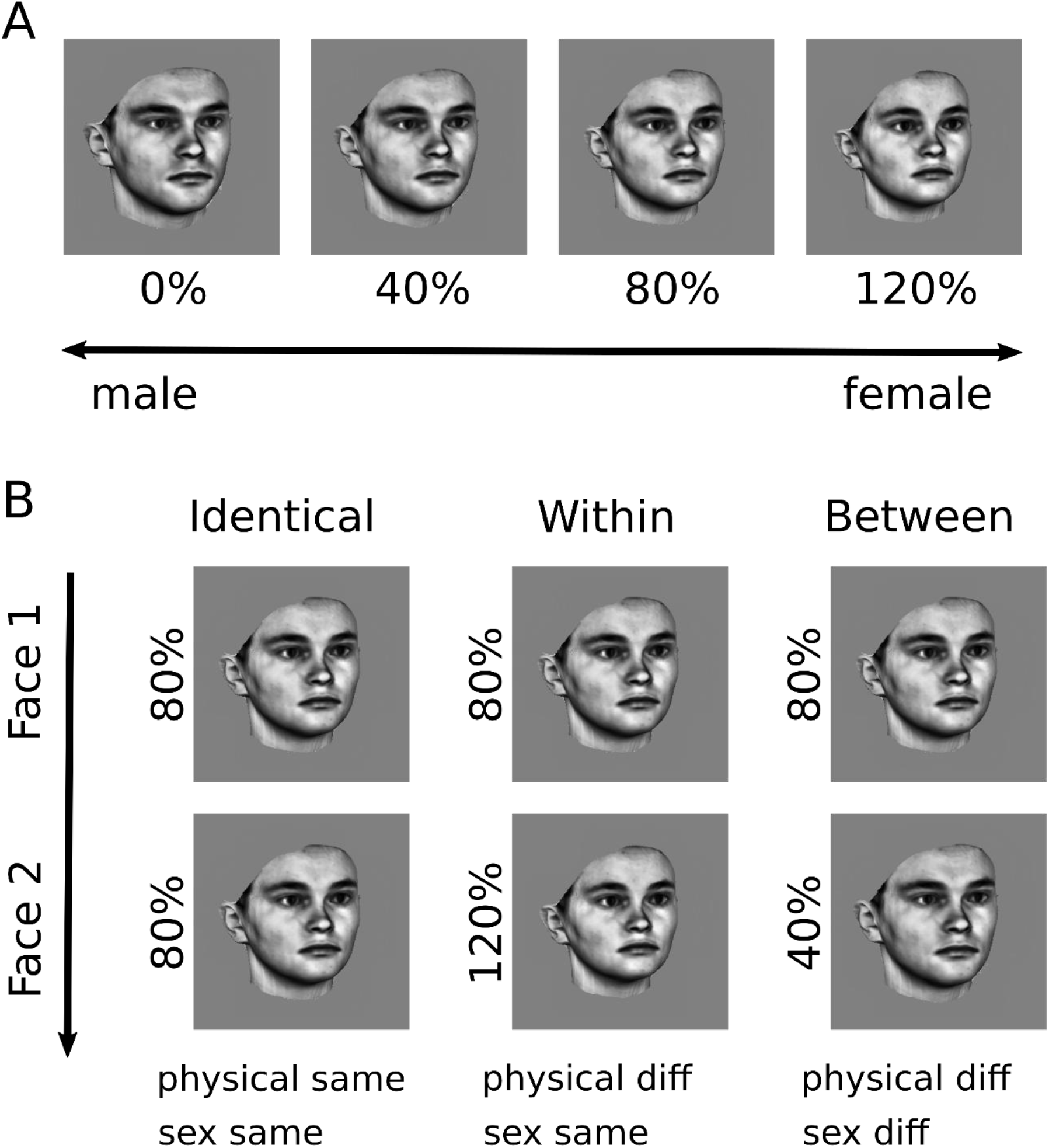
Experimental stimuli and morph conditions. (A) shows the four morph levels of the sex continuum: 0% female (i.e. 100% male), 40% female, 80% female and 120% female. The selected morph levels allowed for the point of subjective equality (change in perception of face sex) to lie between the 40% female and 80% female morph levels, accounting for observers’ male bias in face sex perception (Armann & Bülthoff, 2012). (B) illustrates the three morph conditions for the female continuum direction. In each trial, faces of one identity are shown, and the first face shown in each trial is an 80% female morph. The second face shown in the trial is an 80% female morph for the identical condition, a 120% female morph for the within condition and a 40% female morph for the between condition.

#### 2.2.2. Face localizer stimuli

Stimuli for the face localizer were grayscale images of faces, houses, inverted faces and phase-scrambled images (six exemplars per category). Faces and houses were shown in front of phase-scrambled background images.

### 2.3. Experimental design

#### 2.3.1. Face identity familiarisation procedure

We split our set of 20 face identities into two sets of 10 face identities. Half of our participants were familiarised with one set, and the other half of the participants were familiarised with the other set. The remaining set of faces for each participant group were later used as unfamiliar faces in the fMRI experiment. Participants were familiarised with the male and female endpoint faces of the morph continuum for each identity following the procedure used in Experiment 4 of (Armann & Bülthoff, 2012).

Each participant completed 160 trials in total, where each of the 10 face identities was shown eight times as its male endpoint (i.e. 0% female) and eight times as its female endpoint (i.e. 120% female). Faces were shown in a random order and from varying viewpoints. A question about a character trait was displayed below each face, randomly selected from a list of 46 different traits (e.g. ‘how intelligent is she?’; ‘how attractive is he?’). A male pronoun (i.e. *he*) was used for male endpoint faces and a female pronoun (i.e. *she*) was used for female endpoint faces. Participants responded to each question using a 7-point Likert scale ranging from ‘not very’ to ‘very’.

Following the fMRI experiment, we tested participants’ recognition of the male and female endpoints for all faces (i.e. both familiar and unfamiliar) shown in the fMRI experiment. These 40 faces were shown intermixed with 32 distractor face identities, which were faces that were not shown in the familiarisation procedure or fMRI experiment.

#### 2.3.2. Main experiment procedure (fMRI)

The main experiment consisted of blocks of trials, where each block contained trials from one of six conditions of a 3 (morph conditions) x 2 (familiarity conditions) factorial design. Each participant completed four runs, and in each run five blocks of each of the six conditions were shown, plus five fixation-only blocks, presented in a randomized sequence. Two runs contained stimuli from the female continuum direction (see next paragraph) and two runs contained stimuli from the male continuum direction. Due to an error during data collection, for one participant only two of the four fMRI runs could be used for data analyses (one from each continuum direction).

Each block contained six trials, where each trial consisted of two faces of the same identity shown for 0.5 s one after another, with a 0.5-s grey screen with a fixation cross after each image. For the female continuum direction (see **Fig. 1B**), the first face in each trial was always an 80% female face, and the second face in the trial was an 80% female face for the identical morph condition, a 120% female face for the within morph condition and a 40% female face for the between morph condition. For the male continuum direction, the first face in each trial was always a 40% female face, and the second face in the trial was a 40% female face for the identical morph condition, a 0% female face for the within morph condition and an 80% female face for the between morph condition. The experiment was programmed using Psychtoolbox 3 (Kleiner et al., 2007), http://psychtoolbox.org.

The participants’ task was to detect when a target identity was shown. This identity replaced either the first or the second face (assigned randomly) and was shown in 10% of all trials. The target identity had the same morph level as the face it replaced. Participants made a button press whenever they detected the target identity.

#### 2.3.3. Face localizer procedure (fMRI)

Each participant completed a face localizer immediately following the main experiment. The face localizer consisted of five conditions (faces, houses, inverted faces, phase-scrambled images and fixation cross only), which were presented in a block design. The five conditions were presented in a random sequence. Each block contained six images, where each image was shown for 1 s followed by a 2-s blank grey screen. Participants fixated on a fixation cross throughout the face localizer and performed a one-back matching task: they pressed a button whenever an image was identical to the one preceding it. Repetitions occurred in 5% of trials, with repetition trials assigned randomly.

### 2.4. Imaging parameters

MR images were acquired with a Siemens 3T TIM Trio scanner and a 12-channel phased-array head coil (Siemens, Erlangen, Germany). Functional T2* echoplanar images (EPI) were acquired using a sequence with the following parameters; TR 1.92 s, TE 40 ms, flip angle 90°, FOV 192×192 mm^2^, 27 slices acquired with an interleaved order. Volumes had an in-plane voxel size of 3×3 mm^2^ and slices had a thickness of 3 mm, with a 1-mm gap between slices. The first four volumes of each run were discarded to allow for equilibration of the T1 signal. For each participant, we also recorded a T1-weighted anatomical scan using a sequence with the following parameters; TR 2.3 s, TE 2.98 ms, FOV 240×256 mm^2^, 160 slices, voxel size 1×1×1.1 mm^3^.

### 2.5. fMRI preprocessing

We preprocessed the fMRI data using SPM12 (http://www.fil.ion.ucl.ac.uk/spm/). Functional images were slice time-corrected, realigned and coregistered to the anatomical image. For univariate analyses, images were additionally normalised to MNI (Montreal Neurological Institute) space and smoothed with an 8-mm Gaussian kernel. RSA (representational similarity analyses) were conducted on unsmoothed data in subject space. The resulting searchlight maps were normalised to MNI space and smoothed with an 8-mm Gaussian kernel for the group analyses.

### 2.6. Definition of regions of interest

We defined two face-responsive regions of interest (ROIs) in each hemisphere, the occipital face area (OFA) and the fusiform face area (FFA), using the functional localizer data. **Table 1** shows the mean MNI coordinates and volumes of our OFA and FFA ROIs. We first tried to identify the left and right OFA and FFA for each participant using the contrast faces > houses with a threshold of *p* < .01 uncorrected. For any ROIs we could not define using this contrast, we next tried to identify the ROI using the contrast faces > scrambled images with a threshold of *p* < .01 uncorrected. For each ROI, we selected all voxels within an 8-mm sphere centred on the voxel showing the highest activity. We combined the left and right OFA and FFA components to form one bilateral OFA and one bilateral FFA for each participant.

**Table 1.** Mean MNI coordinates and volumes of OFA and FFA ROIs. Mean MNI coordinates and volumes of our OFA and FFA ROIs (± standard deviations).

| ROI | hem | x |  | y |  | z |  | volume (mm<sup>3</sup>) | N |
| --- | --- | --- | --- | --- | --- | --- | --- | --- | --- |
| OFA | left | -37 | ± 7.2 | -81 | ± 7.2 | -10 | ± 5.5 | 130 ± 120.6 | 12 |
| | right | 42 | ± 5.5 | -76 | ± 6.3 | -11 | ± 6.4 | 227 ± 153.2 | 12 |
| FFA | left | -40 | ± 3.8 | -52 | ± 5.7 | -20 | ± 2.6 | 185 ± 123.8 | 12 |
| | right | 41 | ± 3.7 | -53 | ± 5.6 | -18 | ± 2.1 | 274 ± 163.0 | 12 |

We additionally defined a ROI in the orbitofrontal cortex (OFC) that was previously shown to respond parametrically based on participants’ subjective perception of face sex (Freeman et al., 2010). We transformed the Talairach coordinates given in the paper (−1, 53, -7) to MNI space using the BioImage Suite converter (Lacadie et al., 2008). This gave us MNI coordinates of -1, 59, -5. We selected all voxels in an 8-mm sphere centred on this coordinate to form our OFC ROI.

### 2.7. Behavioural analyses

Participants pressed a button during the fMRI experiment when they detected a target face identity, ensuring they kept their attention on the stimuli. We investigated if there were any differences in the identity detection task performance between our six experimental conditions by performing a 3 (morph condition) x 2 (familiarity condition) repeated measures ANOVA on participants’ percentage correct scores.

### 2.8. fMRI analyses

We modelled each participant’s neural responses with General Linear Models (GLMs) using SPM12. For univariate analyses, GLMs were fitted to normalised and smoothed fMRI data; for RSA, GLMs were fitted to unsmoothed fMRI data in subject space. Each GLM contained regressors for each of the six main conditions, plus a regressor to model the baseline condition, a regressor to model button presses and six realignment regressors that serve as estimates of head motion and were created during the realignment step of the data preprocessing.

#### 2.8.1. Univariate analyses

For ROI analyses, we performed 3 (morph condition) x 2 (familiarity condition) repeated measures ANOVAs to investigate if any ROIs showed differences in blood oxygen level– dependent (BOLD) responses between the six conditions. We tested for non-sphericity using Mauchly’s test of sphericity and, where necessary, applied Greenhouse-Geisser correction for non-sphericity. We also used Bonferroni-correction to adjust for the number of ROIs tested (N = 3). In any cases where we found significant effects of morph condition, or significant interactions between the morph and familiarity conditions, we performed follow-up paired *t*-tests to investigate which conditions showed activation differences. We expected that regions responding to *physical* changes in face sex would show lower activation to the identical morph condition than to the within and between morph conditions and would not show a difference in activation to the within and between morph conditions. In contrast, we expected that regions responding to *categorical* changes in face sex would show a higher activation to the between morph condition than to the within and identical morph conditions. Furthermore, we expected that the *categorical* effect would be stronger for familiar faces, in line with the categorical behavioural judgements of face sex for familiar faces (Armann & Bülthoff, 2012).

For whole-brain analyses, we investigated if any brain regions showed differences in activation that corresponded with BOLD responses to *physical* or *categorical* changes in face sex. To investigate which regions responded to *physical* changes in face sex, we used the contrast: (within morph condition + between morph condition) > identical morph condition. To investigate which regions respond to *categorical* changes in face sex, we used the contrast: between morph condition > (identical morph condition + within morph condition). We performed these analyses separately for unfamiliar and familiar conditions as we predicted that there might be differences in the responses to these conditions for familiar and unfamiliar faces. We assessed group-level significance using nonparametric permutation tests as parametric statistical methods have been demonstrated to be overly conservative for voxel-wise inference, especially for smaller sample sizes such as our N = 12 participants, whereas nonparametric permutation tests have been shown to offer more precise control over the rate of false positives (Eklund et al., 2016; T. E. Nichols & Holmes, 2002; T. Nichols & Hayasaka, 2003). We performed group analyses for each contrast using nonparametric permutation tests with SnPM13 (http://www.nisox.org/Software/SnPM13/), using 8-mm FWHM variance smoothing and 4096 permutations. We assessed significance using voxel-wise inference with a threshold of *p* < .05 and using a family-wise error (FWE) rate correction for multiple comparisons across the whole brain.

#### 2.8.2. Representational similarity analyses (RSA)

We performed RSA analyses to investigate if there were differences in the patterns of activity in any brain region that would be consistent with *physical* or *categorical* neural coding models of face sex. Our *physical* model predicted that patterns of activity would be more similar for the within and between morph conditions than the identical morph condition in regions showing responses to physical changes in face sex. Our *categorical* model predicted that patterns of activity would be more similar for identical and within morph conditions than the between morph condition in regions showing responses to categorical changes in face sex. We conducted all RSA analyses in our three ROIs (transformed into subject-space) and in whole-brain searchlight analyses using 100-voxel spheres. RSA analyses were performed with CoSMoMVPA (Oosterhof et al., 2016) using Tools for NIfTI and ANALYZE image (Shen, 2020). For each ROI and searchlight sphere, we conducted Pearson’s correlations on the pattern of evoked activity between all possible pairs of the identical, within and between morph conditions. We conducted these analyses separately for unfamiliar and familiar faces, and for each fMRI run. We then performed a Pearson’s correlation to compare the differences in similarity between the three morph conditions to dissimilarity matrices consistent with *physical* and *categorical* neural coding models of face sex. We averaged the final correlation values across the four fMRI runs.

For the ROI analyses, as correlation values may not be normally distributed, we performed Wilcoxon signed rank tests to investigate if there was significant correlation between the pattern of activity in each ROI and the physical or categorical models, separately for unfamiliar and familiar faces. We then performed paired Wilcoxon signed rank tests to investigate if there were differences in the correlation with these models between unfamiliar and familiar faces. All ROI analyses were Bonferroni-corrected for the N = 3 ROIs tested.

For searchlight analyses, each participant’s whole-brain correlation maps were first Fisher-transformed to yield normally distributed values. These maps were then normalised to the MNI template and smoothed with an 8-mm Gaussian kernel. We then performed group analyses for each model and each familiarity condition with nonparametric permutation tests in SnPM13 (http://www.nisox.org/Software/SnPM13/), with 8-mm FWHM variance smoothing and 4096 permutations. We assessed significance using voxel-wise inference with a threshold of *p* < .05 and using a FWE rate correction for multiple comparisons across the whole brain.

## 3. Results

### 3.1. Behavioural results

#### 3.1.1. Detection of target identity during the fMRI experiment

Participants performed a target identity detection task during the fMRI experiment to keep their attention on the stimuli. Button presses were not recorded for one participant due to an error during data collection. We performed a 3 (morph condition) x 2 (familiarity condition) repeated measures ANOVA for the remaining 11 participants’ percentage correct scores, in order to investigate if there were any differences in participants’ performance between our six main conditions of interest. Average performance of participants across conditions ranged between 36 and 42 % correct, and we found no significant differences in behavioural performance between any of our conditions (main effect of morph condition: *F*_2,20_ = 1.51, *p* = .24, η_p_^2^ = 0.13; main effect of familiarity condition: *F*_1,10_ = 0.18, *p* = .68, η_p_^2^ = 0.017; interaction between morph and familiarity conditions: *F*_2,20_ = 0.54, *p* = .59, η_p_^2^ = 0.051). Thus, we did not identify any differences in behavioural performance between our experimental conditions.

#### 3.1.2. Recognition of face identities following the fMRI experiment

Following the fMRI experiment, we tested participants’ recognition of the male and female endpoints of the familiar and unfamiliar faces shown in the experiment. On average, participants recognised 71% of familiar faces and 7% of unfamiliar faces, and they showed a false recognition rate of 4% for distractor faces that were not shown in the experiment. The recognition rate for familiar faces was significantly higher than that for unfamiliar faces (*M* = 64%, *SE* = 6.8%, *t*_*11*_ = 9.37, *p* < .001, Cohen’s *d*_*z*_ = 2.71) and for faces that were not shown in the experiment (*M* = 67%, *SE* = 6.4%, *t*_*11*_ = 10.41, *p* < .001, Cohen’s *d*_*z*_ = 3.00). Thus, this result shows that participants were able to recognise the face identities that were shown in the familiarisation procedure significantly better than those that were not. There was no significant difference between the recognition of untrained faces and false recognition of faces that were not shown in the experiment (*M* = 3%, *SE* = 1.7%, *t*_*11*_ = 1.88, *p* = .087, Cohen’s *d*_*z*_ = 0.54), demonstrating that viewing the unfamiliar faces during the fMRI experiment did not lead to familiarity with these faces. We additionally tested whether there were any differences in the recognition of male and female faces and found no significant differences between male and female face recognition for both familiar (*M* = 8%, *SE* = 5.3%, *t*_*11*_ = 1.56, *p* = .147, Cohen’s *d*_*z*_ = 0.45) and unfamiliar (*M* = 1%, *SE* = 1.9%, *t*_*11*_ = 0.43, *p* = .674, Cohen’s *d*_*z*_ = 0.12) faces. This suggests that participants could recognise the female and male faces equally well.

### 3.2. Univariate fMRI results

### 3.2.1. Region of interest results

Figure 2. shows the BOLD responses to the six main conditions in our three ROIs, the OFA, FFA and OFC. We performed 3 (morph condition) x 2 (familiarity condition) repeated measures ANOVAs to investigate if there were differences in BOLD responses between these conditions in any of the ROIs. Both the OFA and FFA showed a significant main effect of morph condition (OFA: *F*_2,22_ = 10.9, *p* < .001 Greenhouse-Geisser-corrected for non-sphericity, η_p_^2^ = 0.50; FFA: *F*_2,22_ = 7.94, *p* = .0025, η_p_^2^ = 0.42), that survived Bonferroni correction for N = 3 ROIs. Neither the OFA nor the FFA showed a main effect of familiarity condition (OFA: *F*_1,11_ = 2.76, *p* = .12, η_p_^2^ = 0.20; FFA: *F*_1,11_ = 2.57, *p* = .14, η_p_^2^ = 0.19) or an interaction between the morph and familiarity conditions (OFA: *F*_2,22_ = 0.71, *p* = .50, η_p_^2^ = 0060; FFA: *F*_2,22_ = 0.016, *p* = .98, η_p_^2^ = 0.0014).

Follow-up paired *t*-tests (**Fig. 2B & 2D**) revealed that both the OFA and FFA showed lower responses to the identical morph condition compared to the within (OFA: *M* = -0.19, *SE* = 0.042, *t*_*11*_ = -4.39, *p* = .0011, Cohen’s *d*_*z*_ = -1.27; FFA: *M* = -0.14, *SE* = 0.035, *t*_*11*_ = -3.97, *p* = .0022, Cohen’s *d*_*z*_ = -1.14) and between (OFA: *M* = -0.086, *SE* = 0.036, *t*_*11*_ = -2.38, *p* = .037, Cohen’s *d*_*z*_ = -0.69; FFA: *M* = -0.098, *SE* = 0.035, *t*_*11*_ = -2.82, *p* = .017, Cohen’s *d*_*z*_ = -0.81) morph conditions. The OFA also showed significantly higher responses to the within than the between morph condition (*M* = 0.10, *SE* = 0.041, *t*_*11*_ = 2.44, *p* = .033, Cohen’s *d*_*z*_ = 0.70), but the FFA showed no significant difference in activation between these two conditions (*M* = 0.041, *SE* = 0.038, *t*_*11*_ = 1.09, *p* = .30, Cohen’s *d*_*z*_ = 0.32). Thus, these results show that both the OFA and FFA are sensitive to *physical* changes in face sex, and that these responses are not affected by face familiarity. Although the OFC showed a slight difference in responses to the unfamiliar and familiar conditions (*F*_1,11_ = 5.74, *p* = .036 Greenhouse-Geisser-corrected for non-sphericity, η_p_^2^ = 0.34), this trend did not survive Bonferroni correction for N = 3 ROIs. The OFC showed no main effect of morph condition (*F*_2,22_ = 0.56, *p* = .57, η_p_^2^= 0.050) or interaction between the morph and familiarity conditions (*F*_2,22_ = 0.25, *p* = .78, η_p_^2^ = 0.022).

**Figure 2.**
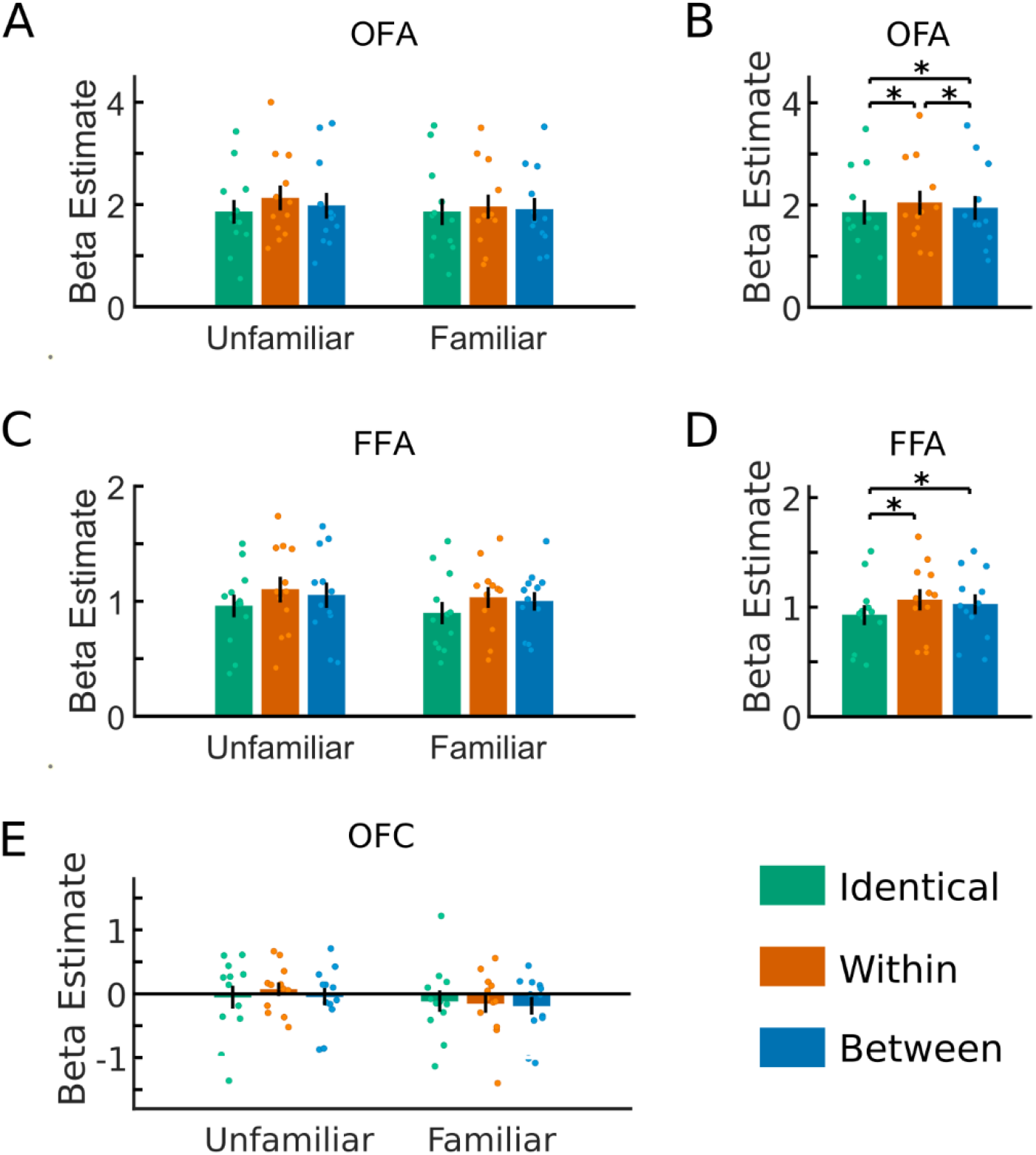
Univariate BOLD responses in the OFA, FFA and OFC. The BOLD responses to all six conditions are shown for the OFA (A), FFA (C) and OFC (E). The BOLD responses to the morph conditions averaged across the two familiarity conditions are shown for the OFA (B) and FFA (D). Bars show the mean response to each condition minus the activation during the baseline fixation condition, and scatter points indicate responses for individual participants. Error bars indicate ±1 SEM. * indicates *p* < .05 in paired *t-*tests.

#### 3.2.2. Whole brain results

We conducted whole brain analyses to investigate if any other regions showed differences in BOLD responses consistent with *physical* responses to face sex (i.e. higher activity to the within and between morph conditions than to the identical morph condition) or consistent with *categorical* responses to face sex (i.e. higher activity to the between morph condition than to the within and identical morph conditions). We conducted group analyses for these two contrasts separately for unfamiliar and familiar faces.

We found significant activation evoked by categorical responses to the sex of familiar faces in one voxel (MNI coordinates: 26, -56, 42) in the right intraparietal sulcus (IPS) (**Fig. 3A** shows this region at a lower threshold of *p* < .1 FWE-corrected). We did not identify any other clusters in any of the other contrasts we tested. We further conducted a 3 (morph condition) x 2 (familiarity condition) repeated measures ANOVA to investigate the BOLD responses to all conditions in the right IPS voxel (**Fig. 3B**). We found a significant main effect of morph condition (*F*_2,22_ = 6.82, *p* = .0050, η_p_^2^ = 0.38) and a significant interaction between the morph and familiarity conditions (*F*_2,22_ = 3.73, *p* = .040, η_p_^2^ = 0.25), but no main effect of familiarity condition (*F*_1,11_ = 0.56, *p* = .47, η_p_^2^ = 0.048).

**Figure 3.**
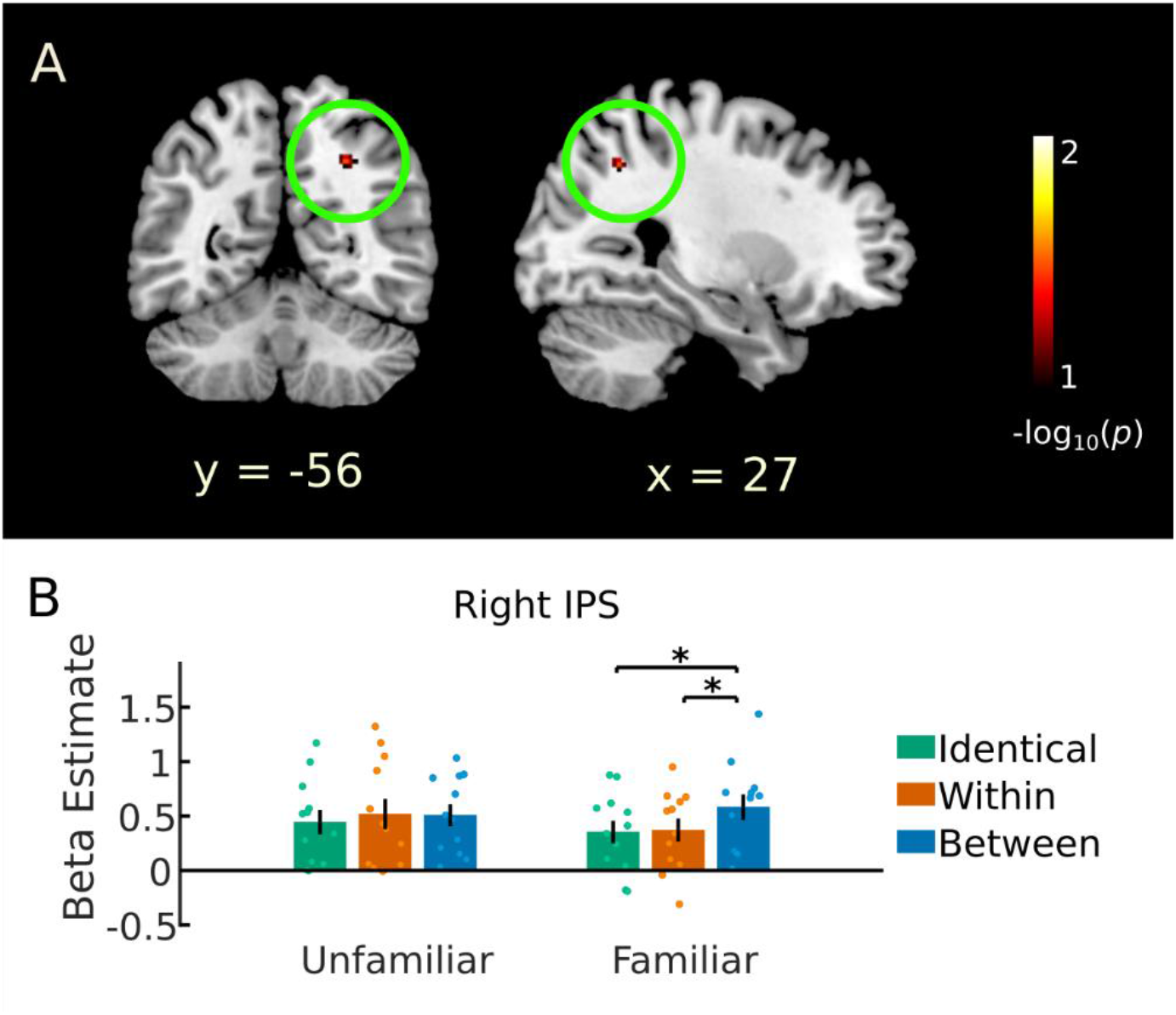
Categorical BOLD responses to face sex in the right intraparietal sulcus (IPS). (A) shows a region in the right IPS (circled in green) showing *categorical* responses to face sex (i.e. stronger activity to the between than within and identical morph conditions) for familiar faces. One voxel in this region was significant at a threshold of *p* < .05 (FWE-corrected); for visualisation, we show the cluster between a lower threshold of –log_10_(*p* values) between 1 (*p* = .1) and 2 (*p* = .01), FWE-corrected. (B) shows the BOLD responses to all six conditions, minus the baseline fixation condition, in the right IPS voxel identified in the whole brain analysis. Bars show mean responses, and scatter points indicate responses for individual participants. Error bars indicate ±1 SEM. * indicates *p* < .05 in paired *t-*tests.

Follow up-paired *t*-tests showed that for familiar faces the right IPS showed significantly higher activity to the between morph condition compared to both the within (*M* = 0.21, *SE* = 0.049, *t*_*11*_ = 4.32, *p* = .0012, Cohen’s *d*_*z*_ = 1.25) and identical (*M* = 0.23, *SE* = 0.050, *t*_*11*_ = 4.59, *p* < .001, Cohen’s *d*_*z*_ = 1.32) morph conditions, but no difference in activation to the within and identical morph conditions (*M* = 0.018, *SE* = 0.054, *t*_*11*_ = 0.33, *p* = .75, Cohen’s *d*_*z*_ = 0.094). For unfamiliar faces, there were no significant differences in activation between any of the three conditions in the right IPS (within and identical: *M* = 0.074, *SE* = 0.049, *t*_*11*_ = 1.53, *p* = .15, Cohen’s *d*_*z*_ = 0.44; between and identical: *M* = 0.064, *SE* = 0.074, *t*_*11*_ = 0.87, *p* = .40, Cohen’s *d*_*z*_ = 0.25; within and between: *M* = 0.010, *SE* = 0.070, *t*_*11*_ = 0.14, *p* = .89, Cohen’s *d*_*z*_ = 0.041). These results show that the BOLD responses in the right IPS followed the expected pattern for categorical coding of face sex for familiar faces, which is consistent with participants’ behavioural judgements of face sex (Armann & Bülthoff, 2012).

### 3.3. Representational similarity analysis (RSA) fMRI results

#### 3.3.1. Region of interest results

Dissimilarity matrices for our three ROIs, showing dissimilarity values that are calculated as 1 – the Pearson’s correlation between pairs of morph conditions, are shown in **Figure 4**. We conducted correlation analyses to investigate if any of our ROIs showed a significant correlation between the differences in BOLD activation patterns to the morph conditions and dissimilarity matrix models consistent with *physical* and *categorical* coding of face sex (**Fig. 5**). We conducted these analyses separately for activity evoked by unfamiliar and familiar faces and conducted Wilcoxon signed rank tests to investigate if any regions showed significant correlations between the dissimilarity models and patterns of BOLD responses. We then performed paired Wilcoxon signed rank tests between the correlations for unfamiliar and familiar faces to investigate if there were any changes evoked by face familiarity.

**Figure 4.**
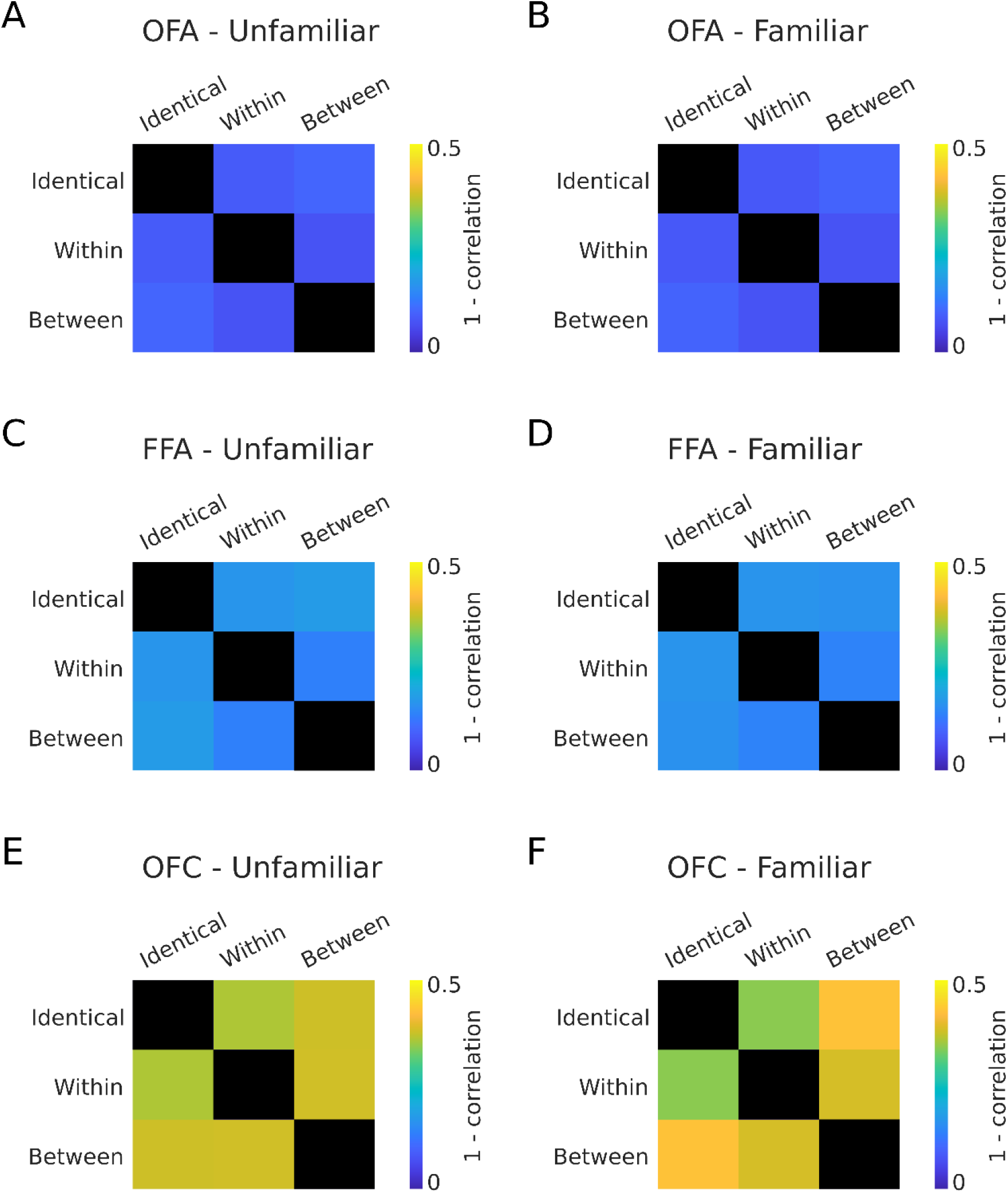
Dissimilarity matrices for unfamiliar (A,C,E) and familiar (B,D,F) faces in the OFA (A,B), FFA (C,D) and OFC (E,F). Dissimilarity colours illustrate 1 – Pearson’s correlation between the patterns of activity evoked by the pairs of morph conditions in each ROI.

**Figure 5.**
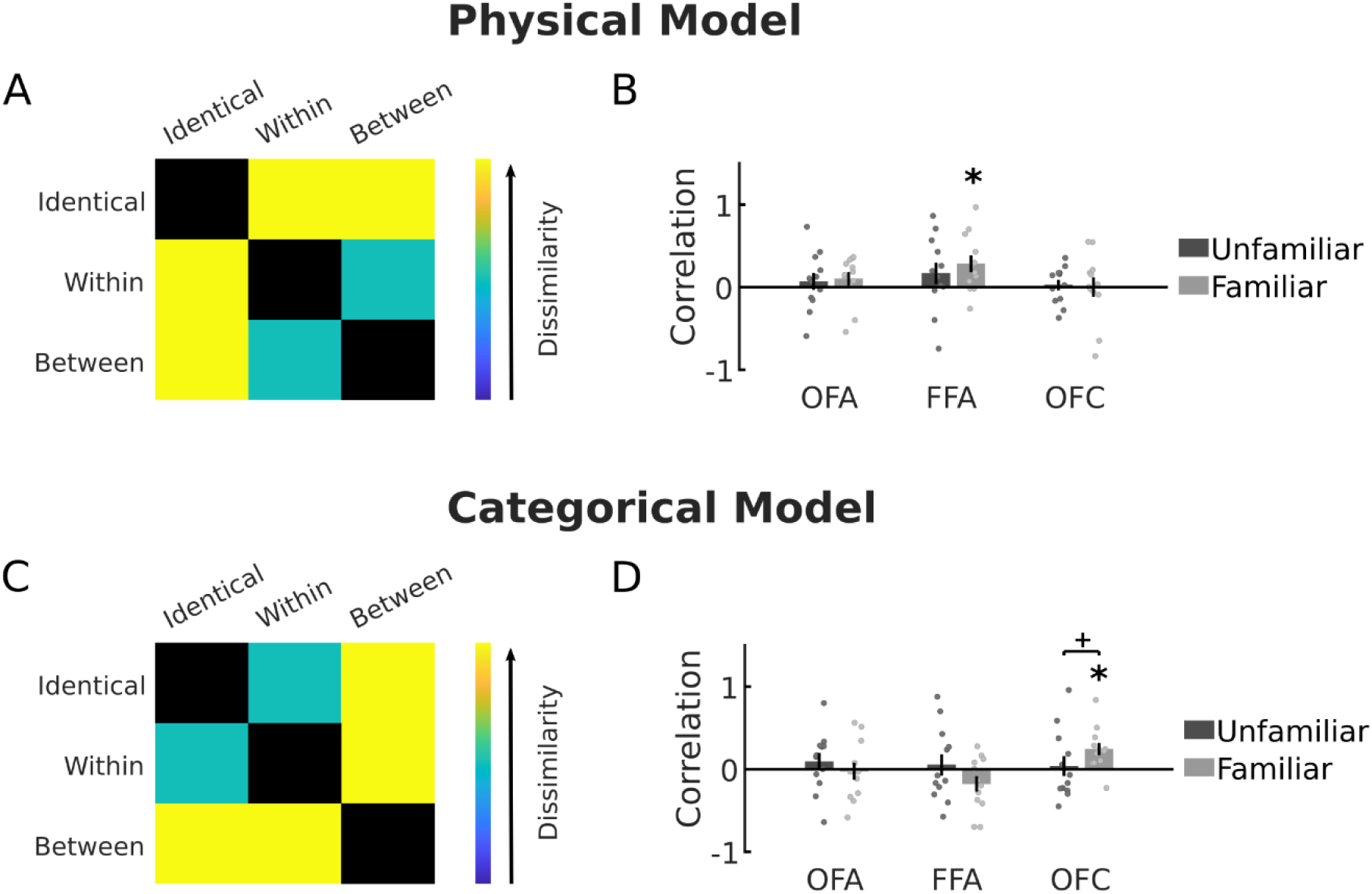
Correlation between ROI dissimilarity matrices (shown in Fig. 4) and the *physical* and *categorical* models of face sex. (A) shows the expected dissimilarity pattern for *physical* coding of face sex, and (B) shows the correlation of this model with the dissimilarity matrices for unfamiliar and familiar faces in the OFA, FFA and OFC. (C) shows the expected dissimilarity pattern for *categorical* coding of face sex, and (D) shows the correlation of this model with the dissimilarity matrices for unfamiliar and familiar faces in the OFA, FFA and OFC. Bars show the mean correlations across participants, scatter points show correlation values for individual participants, and error bars show ±1 SEM. * indicates *p* < .05 in Wilcoxon signed rank tests Bonferroni-corrected for N = 3 ROIs, + indicates *p* < .05 in Wilcoxon signed rank tests uncorrected for N = 3 ROIs.

For the *physical* model (**Fig. 5A**), we found a significant correlation with the activity evoked by familiar faces in the FFA (*median* = 0.23, *Z* = 2.31, *p* = .010), which survived Bonferroni-correction for N = 3 ROIs. We did not find a correlation between the *physical* model and activity evoked by unfamiliar faces in the FFA (*median* = 0.16, *Z* = 1.37, *p* = .085), but we also did not find a significant difference in correlation with the *physical* model between unfamiliar and familiar faces in the FFA (*median* = 0.17, *Z* = 0.71, *p* = .480). We did not find any correlations between the *physical* model and activity in any other ROIs for familiar (OFA: *median* = 0.17, *Z* = 1.22, *p* = .112; OFC: *median* = 0.02, *Z* = 0.35, *p* = .362) or unfamiliar (OFA: *median* = 0.09, *Z* = 0.67, *p* = .253; OFC: *median* = 0.08, *Z* = 0.51, *p* = .305) faces.

For the *categorical* model (**Fig. 5C**), we found a significant correlation with the activity evoked by familiar faces in the OFC (*median* = 0.23, *Z* = 2.55, *p* = .0054), which survived Bonferroni-correction for N = 3 ROIs. We found no correlation of the *categorical* model with activity evoked by unfamiliar faces in the OFC (*median* = -0.12, *Z* = -0.20, *p* = .578); a paired Wilcoxon signed rank test between unfamiliar and familiar correlation with the *categorical* model in the OFC showed a difference in responses (*median* = 0.23, *Z* = 2.04, *p* = .041), but this effect did not survive Bonferroni-correction for N = 3 ROIs. None of the other ROIs showed a significant correlation between the *categorical* model and patterns of activity for familiar (OFA: *median* = -0.03, *Z* = -0.51, *p* = .695; FFA: *median* = -0.17, *Z* = -1.69, *p* = .954) or unfamiliar (OFA: *median* = 0.16, *Z* = 0.98, *p* = .163; FFA: *median* =-0.11, *Z* = 0.27, *p* = .392) faces.

#### 3.3.2. Searchlight results

We performed whole-brain searchlight analyses to investigate if any other brain regions showed differences in the pattern of activity between the morph conditions consistent with *physical* or *categorical* coding models of face sex (separately for unfamiliar or familiar faces). We identified bilateral regions in the medial prefrontal cortex (MPFC) that showed a significant correlation with the *categorical* model for familiar faces (**Fig. 6**). The peak MNI coordinates were -14, 48, 12 for the left hemisphere cluster and 8, 52, 2 for the right hemisphere cluster. For all other contrasts (correlation of unfamiliar faces with the *categorical* model, correlation of unfamiliar faces with the *physical* model and correlation of familiar faces with the *physical* model) we did not identify any significant clusters.

**Figure 6.**
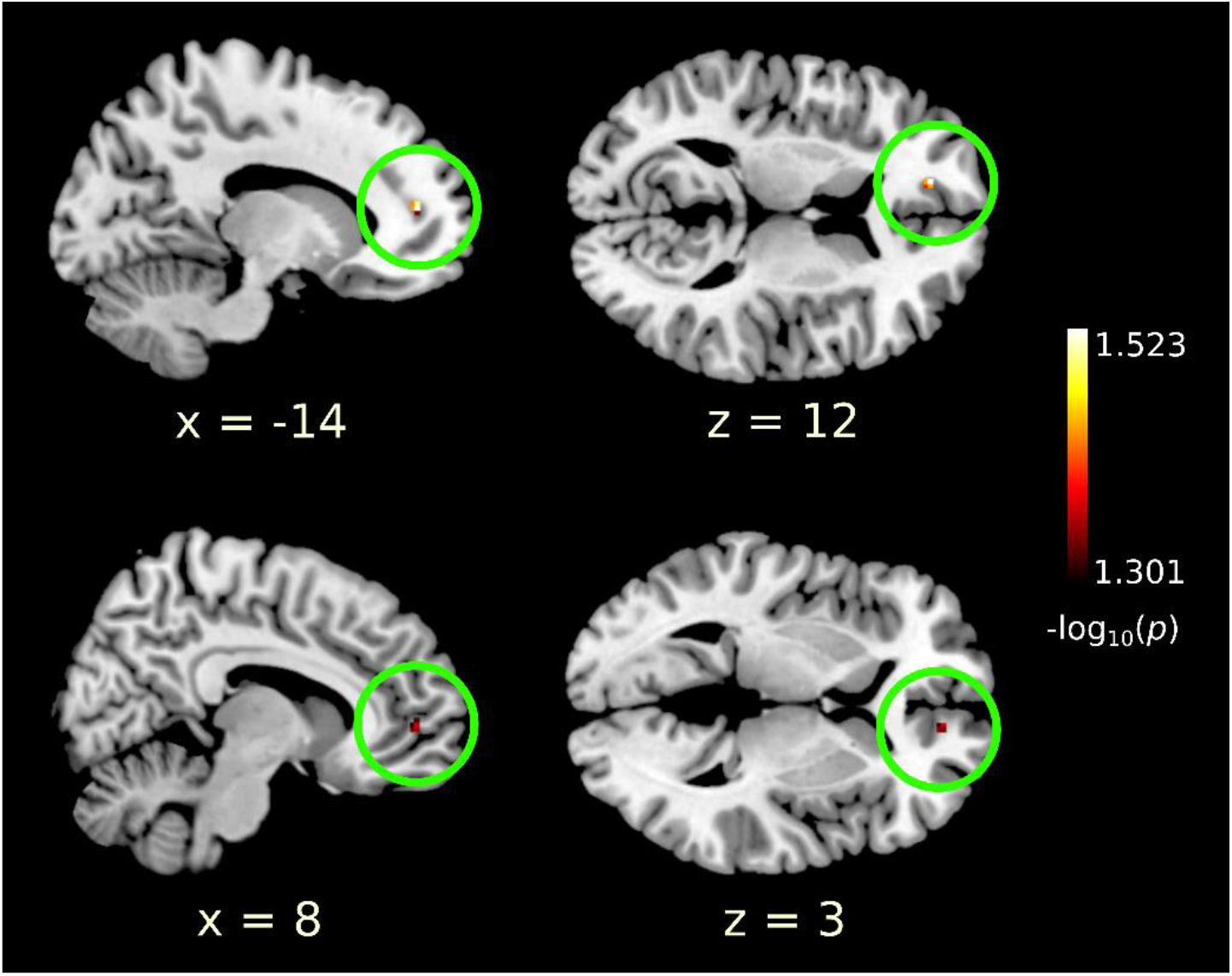
Searchlight results identified bilateral regions in the medial prefrontal cortex (circled in green) that showed a significant correlation of the *categorical* dissimilarity matrix with the patterns of activity evoked by familiar faces. The upper panels show the cluster in the left hemisphere, the lower panels show the cluster in the right hemisphere. –log_10_(*p* values) are shown between 1.301 (*p* = .05) and 1.523 (*p* = .03), FWE-corrected.

## 4. Discussion

In this study, we investigated how the brain encodes the sex of familiar and unfamiliar faces. Participants viewed familiar and unfamiliar faces that were morphed along a continuum between male and female, while their brain activity was recorded using fMRI. Trials showed pairs of faces that either crossed the category boundary between male and female, showed the same amount of morphing but did not cross the category boundary, or were identical. Our results show that the MPFC, OFC and right IPS responded to categorical changes in the sex of familiar, but not unfamiliar, faces. In contrast, the OFA and FFA responded to physical changes in face sex and showed no differences in responses regardless of whether the faces crossed the category boundary or not. Responses in these regions were also unaffected by face familiarity. Altogether, these results show that face-responsive regions in occipitotemporal cortex encode shape-related aspects of face sex, whereas frontal and parietal regions encode learned, categorical aspects of face sex that are linked to face identity.

### 4.1. Categorical coding of the sex of familiar faces

We identified regions in the bilateral MPFC, the OFC and the right IPS that showed categorical coding of the sex of familiar faces. This finding demonstrates that several high-level brain regions show categorical coding of face sex that is linked to face identity. We did not identify any brain regions that showed categorical coding of unfamiliar faces. These results are consistent with behavioural findings that have demonstrated that face sex is perceived categorically for familiar, but not unfamiliar, faces (Armann & Bülthoff, 2012; Bülthoff & Newell, 2004).

Our searchlight RSA analysis identified bilateral regions in the MPFC that showed patterns of responses that correlated with the categorical model of face sex for familiar faces. This finding is compatible with the known roles of the MPFC in social cognition (Amodio & Frith, 2006; Apps et al., 2016). For example, the MPFC is activated when participants recall people they were familiarised with in the context of social situations (Yamawaki et al., 2017), during positive social evaluations of faces (Mende-Siedlecki et al., 2013), and when viewing faces showing incongruent stereotypes (Hehman et al., 2014). Single-cell recordings in the macaque MPFC have revealed neurons that link the identities of others with their specific behaviours during a social decision-making task (Báez-Mendoza et al., 2021). The prefrontal cortex more broadly is known to encode abstract, learned categories (Pan & Sakagami, 2012), and the MPFC has been shown to encode categorical representations of emotional facial expressions in an abstract manner (Murray et al., 2021; Peelen et al., 2010; Skerry & Saxe, 2014), demonstrating a role of this region in categorical coding of faces. Altogether, these findings suggest that the MPFC plays a role in processing and categorising social traits of other people, which is consistent with the present findings of this region encoding the sex of faces linked to familiar identities.

We defined a region in the OFC based on previous coordinates that were found to show responses to face sex in correspondence with participants’ subjective perception of face sex (Freeman et al., 2010). This region has also been proposed to play a general role in encoding social categories of faces in a manner that is biased by participants’ subjective perception (Freeman & Johnson, 2016; Stolier & Freeman, 2016) and neurons in the OFC of macaque monkeys have also been shown to encode social categories of faces (Barat et al., 2018). In our RSA analysis, we found that responses in this region correlated with the categorical model of face sex for familiar, but not unfamiliar, faces. In contrast to our results that are specific to familiar faces, previous studies found that the sex of unfamiliar faces could be decoded from the OFC (Kaul et al., 2011), and that responses in this region were parametrically modulated in line with participants’ subjective perception of the sex of unfamiliar faces (Freeman et al., 2010). However, note that the former study used faces that were uncontrolled for external cues (e.g. hairstyle, make-up) that can often assist sex categorisation, and it is unclear whether the latter study controlled for non sex-related face changes when morphing their face stimuli, which may have led to changes in perceived identity as well as sex. In behaviour, changes in perceived identity that occur alongside changes in perceived sex have been found to allow for categorical perception of unfamiliar faces (Campanella et al., 2001), while unfamiliar faces controlled for identity did not show categorical perception of face sex (Armann & Bülthoff, 2012; Bülthoff & Newell, 2004). Altogether, our results here suggest that categorical coding of face sex in the OFC is either induced or enhanced by face familiarisation.

Our whole-brain univariate analysis identified a region in the right IPS that showed lower responses to pairs of familiar faces with repeated sex than to repeated faces that crossed the sex category boundary. Although the posterior parietal cortex has been classically associated with space, action and attention processing, several studies have shown that it also encodes object categories and identity (Bracci & Op de Beeck, 2016; Vaziri-Pashkam & Xu, 2017, 2019; for a review see Xu, 2018). In particular, the superior IPS was found to encode face identity in an abstract manner (Jeong & Xu, 2016). Our present results show that the right IPS also encodes face sex in a manner linked to face identity. Our coordinates were close to the human hVIP#1, a region proposed to be part of the hVIP complex that encodes face-specific sensory stimuli in relation to their position in external space (Foster et al., 2022). Our present results suggest that this region may also be involved in processing face properties of other people.

### 4.2. Physical coding of face sex

In contrast to the categorical coding of face sex in frontal and parietal regions, two face-responsive occipitotemporal regions, the OFA and FFA, showed responses to physical changes in face sex and were unaffected by face familiarity. Responses in the FFA were lower to repeated identical pairs of faces than to pairs of sex-morphed faces, and showed no differences in responses regardless of whether face pairs crossed the male/female category boundary or not, or were familiar or unfamiliar. A representational similarity analysis showed that response patterns in the FFA were also consistent with physical coding of face sex for familiar faces, and although results were not significant for unfamiliar faces, we did not identify any significant differences in correlations of FFA responses with the physical model for familiar and unfamiliar faces. These results are consistent with previous results showing linear responses in the FFA to sex-morphed faces, in line with the amount of morphing (Freeman et al., 2010), and extend them to show that this coding is not affected by face familiarity. Our present results also highlight a difference between the coding of face sex and face identity in the FFA, as previous work has shown that face identity is encoded categorically in the FFA (Rotshtein et al., 2005). This earlier categorical coding of face identity as compared to face sex may underlie the better precision in recall of identity-specific face features as compared to sex-specific face features (Bülthoff & Zhao, 2020).

The OFA also showed responses consistent with physical coding of face sex, but additionally showed higher responses to face pairs showing a physical change in face sex but not crossing the sex category boundary than to face pairs that did cross the category boundary. We hypothesize that this result might be driven by higher responses evoked by faces at the extremes of the sex-morph continuum, as these faces were present only in the within morph condition, not in the between or identical morph conditions. This result suggests that the OFA may show stronger responses to faces that are more distinct from the average face. Whether the OFA shows such a sensitivity for distinct faces of morph continuums other than sex is unclear. One study that investigated differences in responses to faces morphed in identity did not find higher responses to extreme identity morphs in the OFA; however, they did observe differences between the OFA and FFA in the responses to this condition as compared to identical faces (Rotshtein et al., 2005). These findings are suggestive of differences in how the OFA and FFA code face distinctiveness relative to an average face, however further research comparing the responses in these two regions to faces morphed along different dimensions is needed to fully understand these differences and the underlying face coding.

## Conclusion

In this study, we identify a dissociation between brain regions that encode physical aspects of face sex and categorical face sex aspects that are linked to face identity. We show that face-responsive regions in occipitotemporal cortex respond to physical changes in face sex, but show no effect of changes in sex category or modulation by the familiarity of the viewed faces. In contrast, the bilateral MPFC, OFC and right IPS show categorical responses that are specific to familiar faces, in line with the previously demonstrated specificity of categorical perception of face sex for familiar faces in behaviour.

## Acknowledgements

This research was supported by the Max Planck Society, Germany.

